# Reactive myelopoiesis and FX-expressing monocyte-derived macrophages triggered by chemotherapy promote cancer lung metastasis

**DOI:** 10.1101/2022.12.07.519466

**Authors:** Caijun Wu, Qian Zhong, Rejeena Shrestha, Jingzhi Wang, Xiaoling Hu, Hong Li, Eric C. Rouchka, Jun Yan, Chuanlin Ding

## Abstract

Chemotherapy offers long-term clinical benefits to many cancer patients. However, several pre-clinical studies have demonstrated that certain cytotoxic drugs enhance metastasis via multiple mechanisms. These studies have mainly focused on tumor cell-derived inflammation. The importance of host responses triggered by chemotherapy in regulating cancer metastasis has not been fully explored. Here, we showed that multi-dose Gemcitabine (GEM) treatment promoted breast cancer lung metastasis in a transgenic spontaneous breast cancer animal model. Both CCR2^+^ macrophages and monocytes were increased in the lungs of GEM-treated mice. Further, the increase of CCR2^+^ macrophages and monocytes were observed in naïve (tumor-free) mice after GEM treatment. These changes were largely caused by chemotherapy-induced reactive myelopoiesis that are biased toward monocyte development. Mechanistically, enhanced production of mitochondrial ROS (mtROS) was observed in GEM-treated BM LSK cells and monocytes. Treatment with the mitochondrial targeted antioxidant abrogated GEM induced hyper differentiation of BM progenitors. In addition, GEM treatment induced up-regulation of host cell-derived CCL2, and CCL2/CCR2 axis played essential role in the pro-metastatic host response induced by chemotherapy. Further, GEM and Paclitaxel (PTX) in combination with Doxorubicin (DOX) treatment resulted in up-regulation of coagulation factor X (FX) in lung interstitial macrophages. Targeting activated FX (FXa) using FXa inhibitor or F10 gene knockdown reduced pro-metastatic effect of chemotherapy-triggered host response. Together, these studies suggest a novel mechanism for chemotherapy induced metastasis via the host response-induced accumulation of monocytes/macrophages and interplay between coagulation and inflammation in the lungs.

## Introduction

Metastasis is the primary cause of death in breast cancer patients. The lungs are the second most common site of breast cancer metastasis after the bones (1). Lung metastasis-associated macrophages (MAMs) play very important roles in tumor metastasis through the formation of a pre-metastatic niche (2–6). In mouse models of lung metastasis, interstitial macrophages (IMs, CD11b^high^F4/80^+^CD11c^-^) markedly accumulate in the lungs and differentiate into MAMs (3). Regarding the origin of IMs/MAMs, both tissue-resident macrophages (CCR2^-^) and bone marrow (BM)-derived classical monocytes (BM-Mo, CCR2^+^Ly6C^+^) contribute to the pool of MAMs (7, 8). Several molecules, such as VEGFR1 (6), MMP1 (9), and TGF-β (10), have been identified to participate in the pro-metastatic effects of lung macrophages. However, current strategies targeting macrophage-associated molecules have shown limited success in clinical settings (3).

Chemotherapy offers long-term clinical benefits to many cancer patients. However, several pre-clinical studies have demonstrated that certain cytotoxic drugs enhance metastasis by multiple mechanisms (11–14). These studies have mainly focused on tumor cell-derived cytokines, chemokines, and exosomes (14–17). Our recent studies suggest that Gemcitabine (GEM) promotes the immunosuppressive function of monocytic MDSC (M-MDSC) in the tumor microenvironment via the tumor cell-derived GM-CSF and efferocytosis signaling (18). Emerging evidence indicates that the host response induced by chemotherapy may also play a critical role in regulating tumor progression and metastasis (19). This mechanism may explain why some cancer patient response to chemotherapy over times and tumor re-growth occurs after initially responding to chemotherapy. However, the critical roles of chemotherapy-induced host-responses played in promoting metastasis have not been well understood.

In this study, we showed that GEM treatment promoted breast cancer lung metastasis in a spontaneous breast cancer mouse model with accumulated CCR2^+^ monocytes and macrophages in the lungs. Interestingly, the increase of CCR2^+^ macrophages and monocytes were also observed in tumor-free naïve mice after GEM and combination of PTX and DOX treatment. These changes were largely caused by chemotherapy-induced reactive myelopoiesis, and upregulation of host-cell derived CCL2. In addition, GEM and combination of PTX and DOX treatment resulted in up-regulation of coagulation factor X (FX) in lung macrophages. Inhibition of activated FX (FXa) reduced pro-metastatic effect of host response triggered by chemotherapy. These findings support our hypothesis that host responses triggered by chemotherapy enhance breast cancer lung metastasis via modulation of lung macrophage accumulation and differentiation toward a pro-metastatic phenotype.

## Results

### Gemcitabine chemotherapy promotes breast cancer lung metastasis and accumulation of lung macrophages

Previous studies have shown that certain chemotherapeutic drugs, such as paclitaxel (PTX) (14, 20) and doxorubicin (DOX) (20), promote cancer metastasis in pre-clinical models. We further examined the effects of Gemcitabine (GEM) on lung metastasis because these drugs exhibit different mechanisms of action. MMTV-PyMT mice develop spontaneous mammary tumors that closely resemble the progression and morphology of human breast cancers with poor prognosis (21). The treatment was started at 9-10 weeks of age when the biggest single tumor size reached 6-8 mm in diameter, and last for two weeks. Tumor progression was evaluated for additional 10-14 days after last treatment. Although GEM treatment did not impact the primary tumor progression (Figure 1A), mice received GEM treatment developed more lung metastases than those received PBS control as evidenced by increased tumor nodule counts (Figure 1B). To uncover potential mechanisms underlying pro-metastatic effect of GEM treatment, lung immune cell profile was examined two days later after last GEM treatment using mass cytometry (CyTOF). The cell clustering analysis revealed a significant increase of lung macrophages, particular CCR2^+^ macrophages (Figure 1C). CCR2^+^ macrophages are mainly differentiated from BM-derived monocytes (7, 8). Mass cytometry data also showed an increase of CCR2^+^Ly6C^+^ monocytes in the lungs of GEM-treated mice (Figure 1D). Further, we examined T cell profile in the lungs. GEM treatment increased percentages of naïve CD4 and CD8 T cells (CD44^-^CD62L^+^), whereas effector memory T cells (T_EM_, CD44^+^CD62L^-^) were decreased (Figure1E). Intracellular staining revealed that effector T cells, including IFN-γ-producing CD4/CD8 T cells and granzyme B-expressing CD8 T cells, were significantly decreased in the GEM treated mice (Figure 1F).

**Figure 1.**
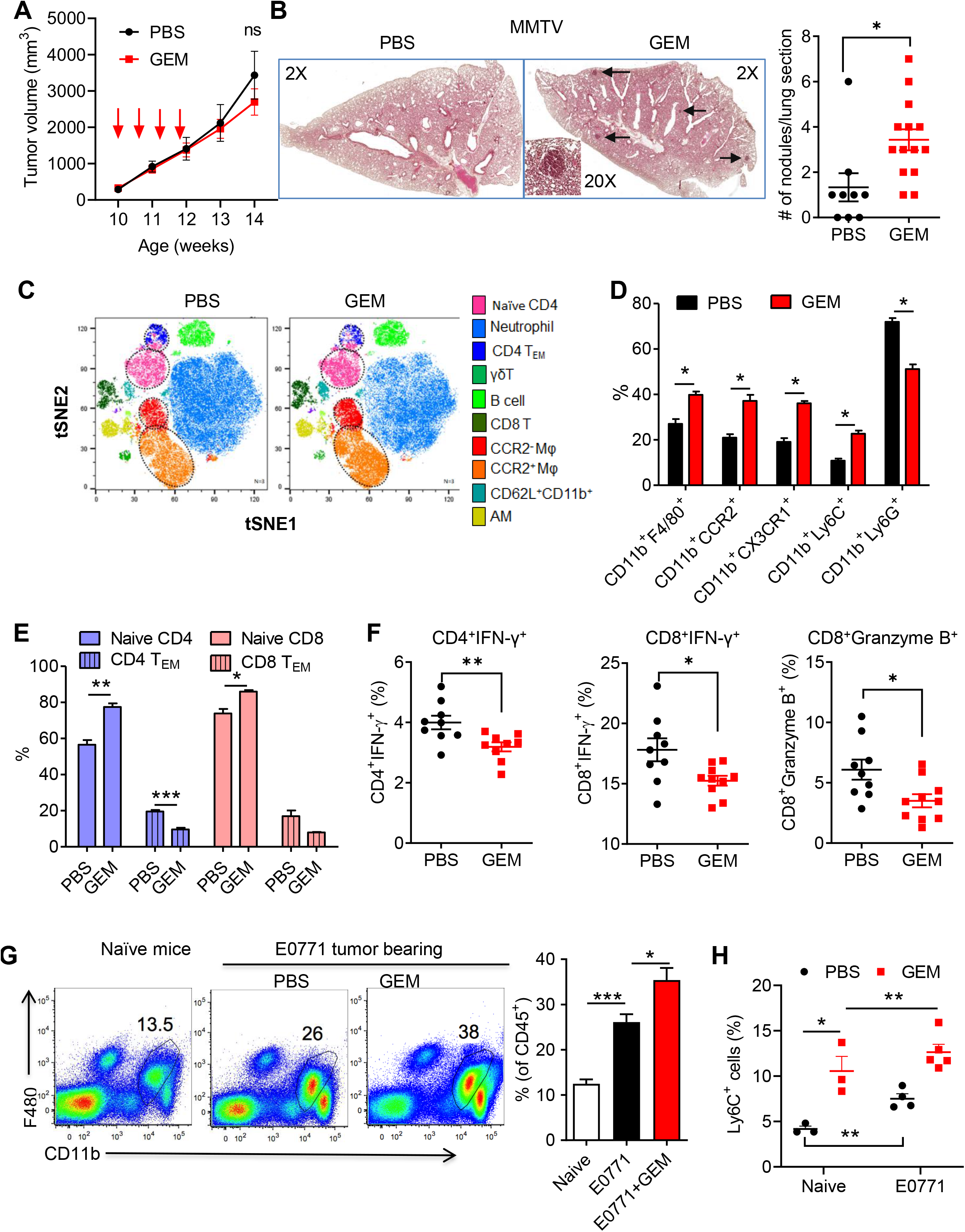
Gemcitabine chemotherapy promotes breast cancer lung metastasis and accumulation of lung macrophages. (**A**) Primary tumor progression after 4 doses of GEM treatment (60 mg/kg, IP). The volume of multiple tumors in each MMTV/PyMT mice was recorded (n=5-6). (**B**) Representative H&E images of lung section of MMTV-PyMT mice from PBS and GEM treated mice. Number of nodules per lung section was summarized (n=9-14). (**C**) Lung tissues were collected two days later. tSNE plot of major immune cell subsets in the lungs of MMTV-PyMT mice identified by FlowSOM clustering algorithm (gated on CD45^+^). (**D**) Summarized data of major myeloid cells from lungs of PBS and GEM treated MMTV-PyMT mice (n=4). (**E**) Summarized data of naïve and effector memory CD4 and CD8 T cells in MMTV-PyMT mice (n=4). (**F**) IFN-γ producing T cells and granzyme B expressing CD8 T cells in MMTV-PyMT mice were evaluated by intracellular staining and flow cytometry (n=9-10). Each dot represents one mouse. (**G**) and (**H**) E077 tumor-bearing mice (tumor size around 6-8 mm in diameter) were treated by 4 doses of GEM in two weeks. Lung macrophages (**G**) and monocytes (**H**) in naïve and tumor-bearing mice were analyzed by flow cytometry (n=3-5). Data are representative of two or three independent experiments and presented as mean ± SEM. *ns*= not significant, \**p* <0.05, \*\**p* <0.01, \*\*\**p* <0.001.

We further addressed the effects of chemotherapy on myeloid cells in the lungs using E0771 tumor-bearing mice. E0771 is characterized between luminal B and triple negative subtype and sensitive to various cancer therapy (22). Our previous studies have shown that multi-dose of GEM treatment results in a reduction of primary tumor progression but increase of Ly6C^+^ myeloid cells in primary tumor microenvironment (TME) (18). We further examined the effects of chemotherapy on lung myeloid cells and found that GEM treatment significantly increased the accumulation of lung MAM (CD11b^high^F4/80^+^CD11c^-^) and monocytes (CD11b^high^Ly6C^+^Ly6G^-^) in E0771 tumor-bearing mice after GEM treatment (Figure 1G and H). These data suggest that host responses triggered by chemotherapy might contribute to modulation of myeloid cells in lung tissue. We further characterized and compared T cells presented in the lungs of chemotherapy treated mice. The remarkable decrease of IFN-γ-producing CD4 and CD8 T cells was observed in the lungs of E0771 tumor-bearing mice received GEM treatment (Supplemental Figure 1). Together, these data suggest that certain chemotherapy treatment may promote lung metastasis through recruitment of CCR2^+^ macrophages and monocytes into the lungs and subsequent impaired T cell function.

### Chemotherapy-triggered host responses promote tumor metastasis in mice

The roles of tumor cell-derived cytokines, chemokines, and exosomes in chemotherapy-induced metastasis have been previously investigated (13, 14). To examine whether host-responses following chemotherapy also modulate lung myeloid cells, tumor-free naïve mice were treated with GEM. Similar to tumor-bearing mice, GEM treatment induced accumulation of macrophages (CCR2^+^CD11b^+^F4/80^+^) and monocytes in the lungs (CCR2^+^Ly6C^+^) (Figure 2A and B). CCR2^+^ macrophages are replenished through monocyte recruitment. Thus, we further examined monocytes in the BM and found that more monocytes were generated in the BM of GEM-treated mice (Figure 2C). A BrdU incorporation assay revealed enhanced proliferation of Ly6C^+^ cells in GEM-treated mice (Figure 2D), as BrdU is only incorporated into newly synthesized DNA of proliferating cells. More importantly, BM monocytes from GEM-treated mice displayed immunosuppressive feature when co-culture with OVA-TCR transgenic T cells (Figure 2E).

**Figure 2.**
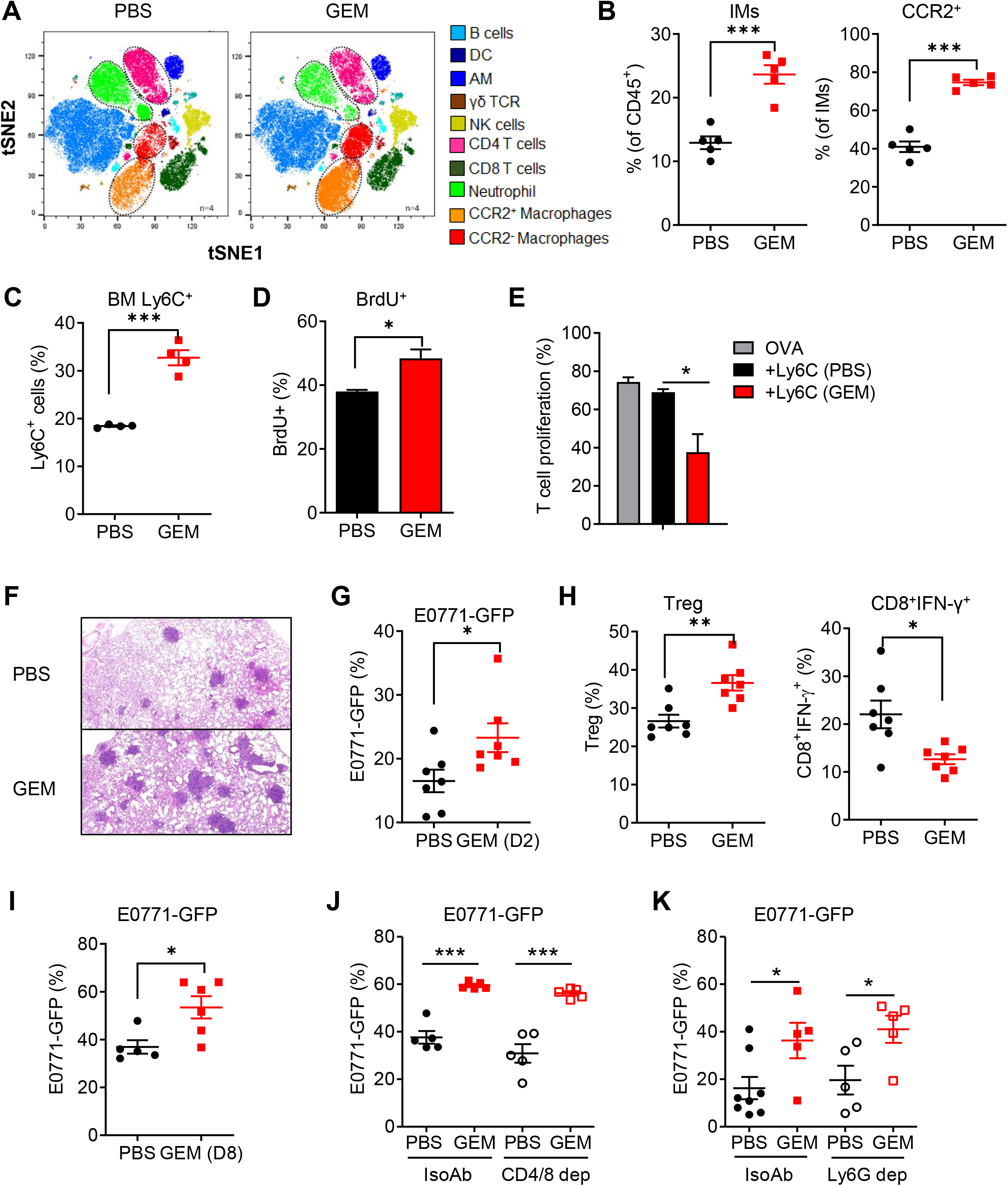
Chemotherapy-triggered host responses promote tumor metastasis in mice. Naïve tumor-free C57Bl/6 mice were treated with 4 doses of GEM (60 mg/kg, IP). Lung tissues and bone marrow were harvested two days later after last treatment. (**A**) tSNE plot of major immune cell subsets in the lungs of PBS and GEM treated mice identified by FlowSOM clustering algorithm (gated on CD45^+^). (**B**) Summarized data of lung macrophages (CD11b^high^F4/80^+^CD11c^-^) and CCR2^+^ macrophages from PBS and GEM treated mice (n=5). (**C**) Summarized data of BM Ly6C^+^ monocytes gated on CD11b^+^ cell population (n=4). (**D**) The PBS and GEM treated mice were intraperitoneally injected with BrdU (2 mg per mouse). BM was collected 16 hours later and the proliferating Ly6C^+^ cells stained for incorporated BrdU were analyzed by intracellular staining and flow cytometry (n=3). (**E**) BM Ly6C^+^ cells from PBS and GEM treated mice were sorted and co-cultured with CFSE-labelled OT-I splenocytes (1:1 ratio) in the presence of OVA (20 μg/ml) for 3 days. T cell proliferation was measured by flow cytometry (n=4). (**F**) Tumor-free mice were treated with 4 doses of GEM and PBS, followed by intravenous injection of E0771-GFP cells (4×10^5^ per mouse) two days later after last GEM treatment. Lung metastasis was determined at day 14 by histological analysis. (**G**) Tumor burden in lungs of GEM pre-treated mice was determined by measuring GFP^+^ tumor cells within CD45^-^ cell population (n=7). (**H**) Regulatory T cells and IFN-γ producing CD8 T in GEM pre-treated tumor-bearing mice were evaluated by intracellular staining and flow cytometry (n=7). (**I**) E0771-GFP cells (4×10^5^ per mouse) were IV injected into GEM pre-treated mice eight days later after last GEM treatment. Lung metastasis was determined by measuring GFP^+^ tumor cells within CD45^-^ cell population (n=5-6). (**J**) and (**K**) B6 tumor-free mice were treated by 4 doses of GEM and PBS. The Ly6G depletion Ab (300 μg, IP, twice a week) (**J**) or CD4/CD8 depletion Abs (250 μg, IP, weekly) (**K**) and IsoAb were used during the GEM or PBS pre-treatment. The experimental lung metastasis assay was performed as described in Figure 3F, and lung metastasis was determined by measuring GFP^+^ tumor cells within CD45^-^ cell population (n=5-8). Data are representative of two independent experiments and presented as mean ± SEM. Each dot represents one mouse. \**p* <0.05, \*\**p* <0.01, \*\*\**p* <0.001.

To investigate the consequence of host-specific responses following chemotherapy on metastasis, naïve mice were treated with four times of GEM in two weeks, followed by intravenous injection of E0771-GFP cells. Lung metastasis was significantly higher in the GEM pre-treated mice compared to that in PBS treated control mice, as determined by histopathological analysis with routine H&E staining (Figure 2F) and flow cytometric analysis of GFP^+^ tumor cells (Figure 2G). The effector T cells (CD8^+^IFN-γ^+^) were decreased, whereas Treg cells were increased in the GEM-pretreated mice (Figure 2H). We further performed experiment by injection of tumor cells 8 days later after chemotherapy when BM HSPCs recover from chemotherapy-induced stress (23). Lung metastasis was also higher in the GEM pre-treated mice compared to that in PBS treated control mice (Figure 2I). These data support the importance of host-response in chemotherapy-induced lung metastasis.

Chemotherapy has been shown to modulate T cell compartment and function in breast cancer patients (24). Previous studies also revealed that Ly6G^+^ neutrophils support lung colonization of metastatic cancer cells (25). To test whether T cells and neutrophils are critical for the pro-metastatic effect triggered by chemotherapy, CD4/CD8 and Ly6G depletion antibody were used during the GEM or PBS treatment period. No difference of lung tumor burdens was observed in the IsoAb and T cell or neutrophil depleted GEM pre-treated mice (Figure 2J and K). These data demonstrate the importance of monocytes and BM-derived macrophages, but not T cells and neutrophils, in pro-metastatic effects of host-response triggered by chemotherapy.

### GEM treatment induces reactive myelopoiesis with enhanced myeloid potential

Lung macrophages include tissue-resident macrophages and BM-derived macrophages, which are differentiated from CCR2^+^ classical monocytes. We hypothesized that the elevated frequency of monocytes/macrophages in post-GEM treatment may arise from an increased number of BM myeloid progenitor cells that generate more BM monocytes. To determine whether GEM treatment induces reactive myelopoiesis, naïve and E0771 tumor-bearing mice were treated by four times of GEM. E0771 primary tumor development induced an increase of BM Lin^-^Sca1^+^c-Kit^-^ cells (LSKs) and multipotent progenitors (MMPs, CD48^+^CD150^-^). GEM treatment further increased the accumulation of BM LSKs and MPPs (Figure 3A and B).

**Figure 3.**
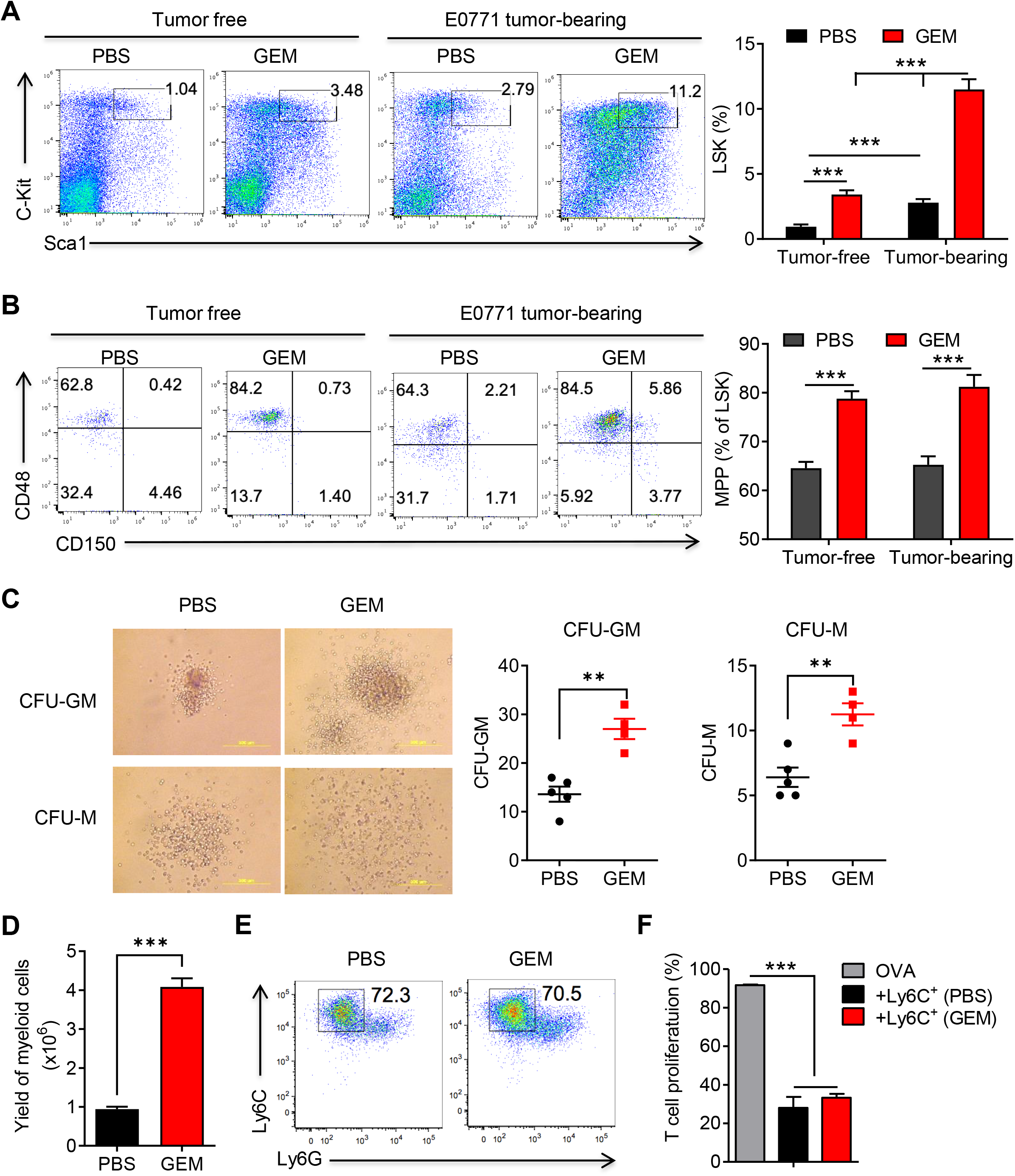
GEM treatment induces reactive myelopoiesis in tumor-free and tumor-bearing mice with enhanced myeloid potential. Naïve C57Bl/6 and E0771 tumor-bearing mice were treated with 4 doses of GEM (60 mg/kg, IP). Bone marrow was harvested two days later after last treatment. (**A**) Representative FACS plots gated on lineage negative cells and summarized data of LSK cells (Lin^-^Sca-1^+^c-Kit^+^) (n=5-8). (**B**) Representative FACS plots gated on LSK cells and summarized data of MPPs (CD150^-^CD48^+^) (n=5-8). (**C**) Representative CFU-GM and CFU-M after 7 days culture of BM cells from GEM-treated tumor free mice in MethoCult™ GF M3534 methylcellulose-based medium. Numbers of CFU-GM and CFU-M were summarized (n=4-5). (**D**) BM cells (1×10^6^) were cultured in the presence of 20% E0771 conditioned medium (CM) for 2 days. The culture medium including non-adherent cells was entirely discarded at day 3 and replaced by medium containing E0771 CM for additional four days. The yield of myeloid cells wase counted after 6 days culture (n=3). (**E**) Representative FACS plots of Ly6G^-^Ly6C^+^ *in vitro* expanded cells. (**F**) Sorted Ly6G^-^Ly6C^+^ cells were cultured with CFSE-labeled OT-I splenocytes in the presence of OVA (20 μg/ml) for 3 days. T cell proliferation was measured by flow cytometry (n=3). Data are representative of two or three independent experiments and presented as mean ± SEM. \*\**p* <0.01, \*\*\**p* <0.001.

To examine the ability of BM progenitors to differentiate into monocytes and macrophages, we performed the colony-formation assay using BM cells from GEM- or PBS-treated mice. After 7 days of culture, both colony numbers of CFU-GM and CFU-M from GEM-treated BM cells were increased as compared to that from PBS-treated BM cells (Figure 3C). No changes of CFU-G were observed between two groups. We further cultured BM cells from GEM or PBS treated mice in the presence of E0771 tumor cell conditioned medium (CM). There was a significant increase in the yield of cells from GEM-treated BM cells after 6 days *in vitro* culture (Figure 3D). The major cell population displayed the phenotype of monocytes (CD11b^+^Ly6C^+^Ly6G^-^) (Figure 3E). Importantly, these *in vitro* differentiated cells exhibited potent immunosuppressive function when co-culture with T cells (Figure 3F). Together, these data suggest that multi-dose chemotherapy may induce reactive myelopoiesis with myelopoietic bias that boost monocyte development and expansion, ultimately leading to the accumulation of immunosuppressive monocyte-derived macrophages in the lungs.

### Up-regulation of mitochondrial ROS (mtROS) in BM microenvironment triggered by GEM treatment

In response to various types of inflammation, HSPCs undergo a metabolic switch from glycolysis to oxidative phosphorylation (OXPHOS) that leads to increased ROS production, which is important for HSPC proliferation and differentiation (26–30). We observed a significant increase of mtROS in LSK cells (Figure 4A) and monocytes (Figure 4B) from mice received GEM treatment. Mitochondria that have a high membrane potential are more prone to ROS generation (31). Thus, we measured mitochondrial membrane potential by using TMRM staining, and found higher mitochondrial potential in LSK cells of GEM-treated mice (Figure 4C). Mitochondrial dysfunction is associated with increased ROS production (32). We further stained BM cells with MitoTracker Green and MitoTracker Red to distinguish between functional mitochondria (MitoTracker Red^high^) and dysfunctional mitochondria (MitoTracker Green^high^, MitoTracker Red^+/low^) (33). An increase of dysfunctional mitochondria but decrease in functional mitochondria in GEM-treated LSKs compared with those in control mice was observed (Figure 4D). These data suggest that GEM treatment has a potential to modulate HSPC mitochondrial activity and metabolism.

**Figure 4.**
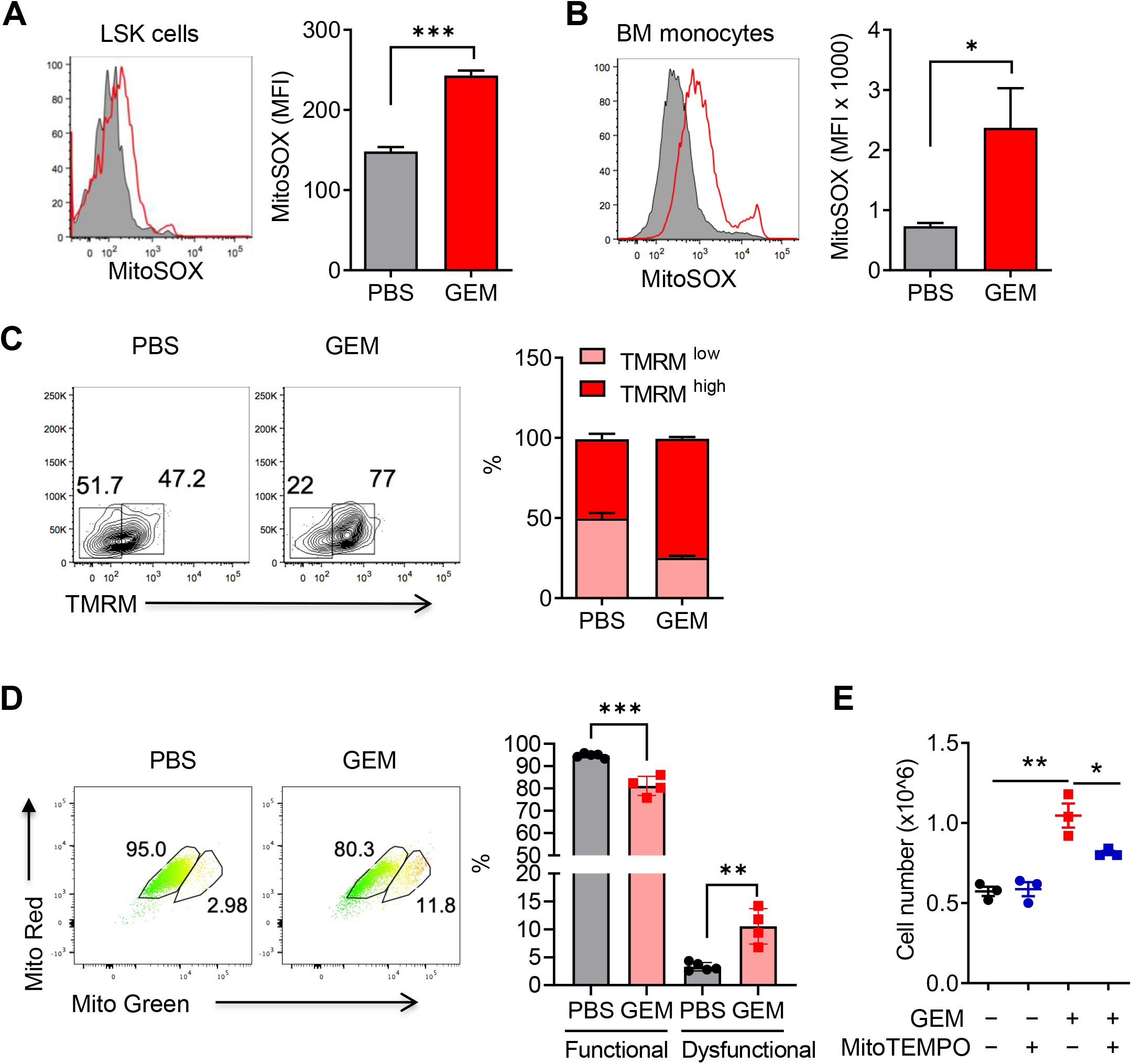
Up-regulation of mtROS in BM microenvironment triggered by GEM treatment. Naïve tumor-free C57Bl/6 mice were treated with 4 doses of GEM (60 mg/kg, IP). Bone marrow was harvested two days later after the last treatment. (**A** and **B**) mtROS production in LSKs (**A**) and BM monocytes (**B**) was determined by using dye MitoSOX (5 μM) (n=3-5). (**C**) Mitochondria potential was determined by using dye TMRM (500 nM) followed by lineage markers and Sca-1, c-Kit Abs staining. The percentages of TMRM^high^ and TMRM^low^ LSK cells were summarized (n=3-4). (**D**) BM cells were stained with MitoTracker Green and MitoTracker Red followed by BM LSK cell markers. Representative plots gated on LSKs and summarized functional and dysfunctional mitochondria were shown (n=4). (**E**) BM cells (1×10^6^) from PBS and GEM pre-treated mice were treated with MitoTEMPO (10 μM) and cultured in the presence of GM-CSF (20 ng/ml) for 2 days. The culture medium including non-adherent cells was entirely discarded at day 3 and replaced by medium containing GM-CSF and MitoTEMPO for additional four days. Cell numbers of CD11 b^+^Ly6C^+^ monocytes after 6 days culture was determined by cell counting and flow cytometry (n=3). Data are representative of two experiments and presented as mean ± SEM. \**p* <0.05, \*\**p* <0.01, \*\*\**p* <0.001.

Emerging studies suggest that ROS also acts as signal-transducing molecules that drive hematopoietic stem cell self-renewal and emergency granulopoiesis (34). To determine the importance of mtROS in GEM treatment induced myelopoiesis, BM cells from GEM and control mice were treated with mitochondria-targeted superoxide scavenger mitoTEMPO and then cultured in the presence of GM-CSF (35). Significant increase of *in vitro* differentiated monocytes was observed from the BM of GEM-treated mice, and inhibition of mtROS using mitoTEMPO abrogated GEM treatment induced high yield of monocytes (Figure 4E). These results suggest that mtROS production in BM niche might play important role in chemotherapy-induced BM myelopoiesis and monocyte development.

### Chemotherapy-induced up-regulation of macrophage-synthesized FX contributes to lung metastasis

To identify factors that may contribute to the phenotypic changes of lung macrophages after GEM treatment, we performed an unbiased gene expression profiling analysis of macrophages from GEM and PBS-treated mice. A total of 2712 differentially expressed genes (DEGs) were recorded (1315 upregulated DEGs and 1397 downregulated DEGs). Coagulation factor X (F10) was identified as one of the most upregulated differentially expressed genes in the macrophages from GEM-treated mice (Figure 5A), which was validated by using qRT-PCR (Figure 5C). Gene ontology (GO) analysis revealed a pattern of enrichment in pathways related to leukocyte adhesion and migration, negative regulation immune cell process, as well as regulation of vasculature development (Figure 5B). Additionally, levels of total FX and FXa were significantly increased in the plasma of mice received GEM treatment (Figure 5D). Further, several pro-metastatic molecules, including S100A8, S100A9, and TGF-β, were up-regulated in the lung tissues after GEM treatment (Supplemental Figure 2).

**Figure 5.**
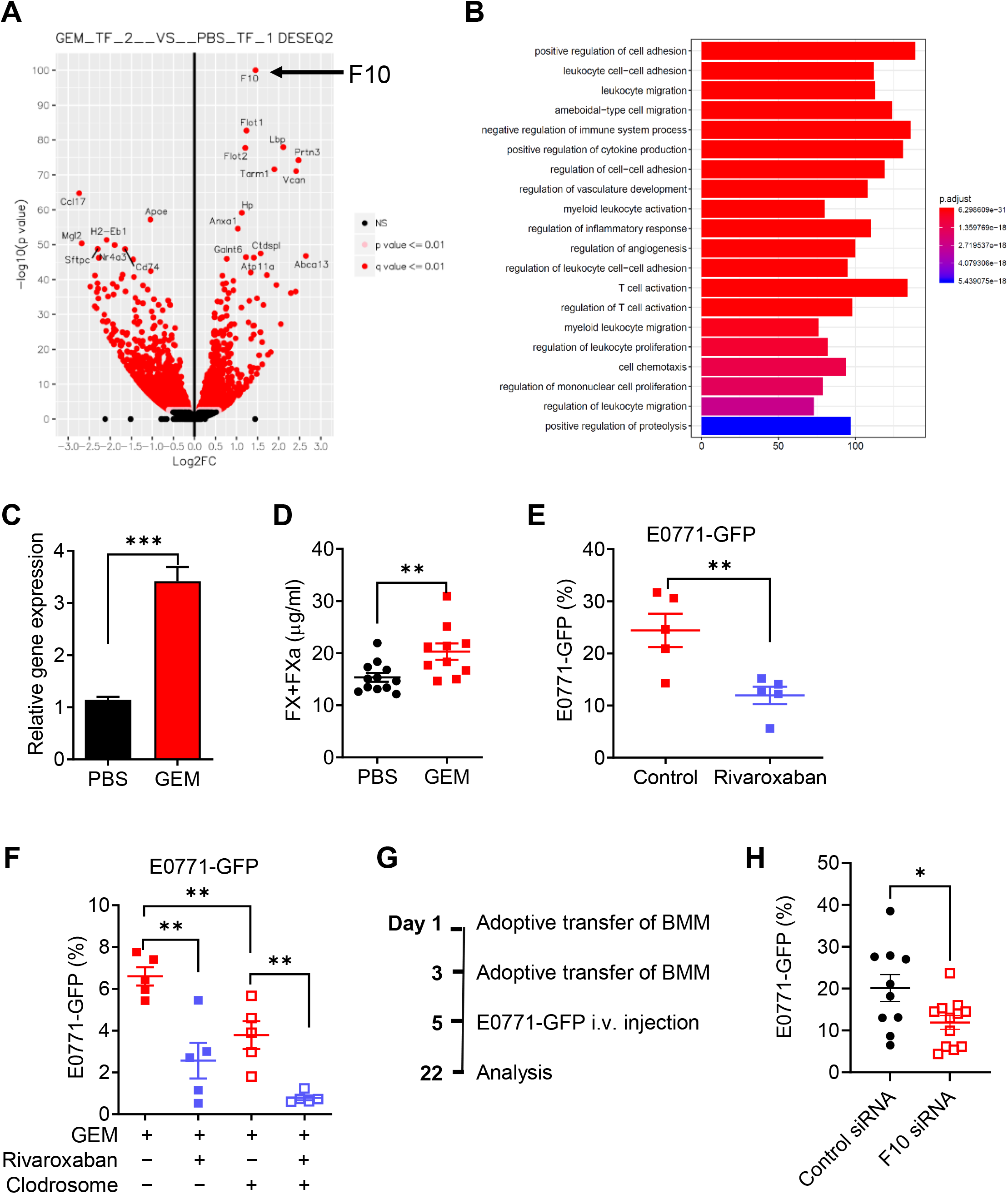
Chemotherapy-induced up-regulation of macrophage-synthesized FX contributes to lung metastasis. (**A**) RNA-seq analysis in lung macrophages sorted from naïve mice after GEM and PBS treatment (n=3). Volcano plot for differential gene expression in lung macrophages from GEM-treated mice as compared to PBS-treated mice. (**B**) Top 20 enriched GO biological process for the genes in lung macrophages from GEM-treated mice, as compared to PBS-treated mice. (**C**) qRT-PCR for F10 gene in lung macrophages from GEM and PBS treated mice. (**D**) Levels of total FX and FXa in the plasma of GEM and PBS treated mice (n=10-12). (**E**) Naïve mice were treated with GEM and Rivaroxaban, or solvent control followed by IV injection of 4×10^5^ E0771-GFP cells. The lung tumor burden was examined by measuring GFP^+^ tumor cells within lung CD45^-^ cells 14 days later after tumor cell injection (n=5). (**F**) Mice were treated with GEM and Rivaroxaban, or solvent control followed by IV injection of 1×10^5^ E0771-GFP cells. Lung macrophages were depleted by IV injection of Clodrosome during the GEM/Rivaroxaban treatment and tumor development. The lung tumor burden was examined by measuring GFP^+^ tumor cells within lung CD45^-^ cells 14 days later after tumor cell injection (n=5). (**G**) Experimental scheme for adoptive transfer of E0771 CM stimulated (24 h) control and F10 gene knockdown macrophages (2×10^6^/mouse) and E0771-GFP (4×10^5^/mouse) injection. (**H**) The lung tumor burden was examined by measuring GFP^+^ tumor cells within lung CD45^-^ cells 17 days later after tumor cell injection (n=10-12). The data in panel **H** represents a combination of two experiment. Each dot represents one mouse. \**p* <0.05, \*\**p* <0.01.

To determine the role of macrophage expressed FX in the GEM induced pro-metastatic effect *in vivo,* mice were pre-treated with GEM along with Rivaroxaban, an orally inhibitor of activated form of FX (FXa), and vehicle control daily throughout the GEM pre-treatment period. E0771-GFP cells were intravenously injected two days later after the last GEM treatment. As shown in Figure 5E, inhibition of FXa using FXa inhibitor Rivaroxaban significantly reduced pro-metastatic effect of host-response triggered by GEM treatment. To address whether the anti-metastatic effect of FXa inhibitor is mediated by macrophage expressed FX, we depleted lung macrophages by intravenous injection of Clodrosome. The experimental scheme for *in vivo* GEM and Rivaroxaban treatment and macrophage depletion efficacy were shown in Supplemental Figure 3. Depletion of macrophages resulted in reduced tumor burden in the GEM pre-treated mice. The anti-metastatic effect of FXa inhibition was also observed in the macrophage-depleted mice (Figure 5F). These data suggest that lung macrophages are critical for the pro-metastatic effect of chemotherapy. The off-target effect of FXa inhibitor may exist.

To further determine the roles of macrophage specific FX in the pro-metastatic effect of chemotherapy, we performed *in vitro* F10 gene knockdown in BM-derived macrophages using F10 siRNA. The knockdown efficiency was shown in Supplemental Figure 4. The E0771 CM treated F10 knockdown and control macrophages were adoptively transferred into CCR2 KO recipient mice twice with 48 hours apart, followed by iv injection of E0771-GFP cells (Figure 5G). As shown in Figure 5H, the mice transferred with F10 knockdown macrophages had lower percentage of GFP^+^ tumor cells compared to mice transferred with control macrophages, indicating the contribution of macrophage expressed FX in the pro-metastatic effects of chemotherapy.

Intravenous cancer cell injection does not recapitulate all the steps of metastasis. We performed experiments to further evaluate the anti-metastatic effect of Rivaroxaban in E0771-GFP subcutaneous tumor model. E0771 is poorly metastatic as compared to 4T1 tumors. We have performed three rounds of *in vivo* passage. E0771-GFP cells recovered from tumor-bearing lungs can develop pulmonary metastases. Mice bearing E0771-GFP subcutaneous tumors were treated with Rivaroxaban (oral, 20 mg/kg, daily) or solvent control for two weeks. As shown in Supplemental Figure 5, Rivaroxaban treatment significantly decreased lung metastasis, which was observed in 3 out of 14 Rivaroxaban treated mice compared to 9 out of 14 control mice. The mechanism underlying the anti-metastatic effect of FXa inhibitor in primary tumor-bearing mice need to be further defined.

### GEM-induced pro-metastatic host response is dependent on CCL2/CCR2 mediated recruitment of monocytes

Primary tumor-derived CCL2 can enhance breast cancer metastasis assisted by the recruitment of Ly6C^+^CCR2^+^ monocytes (36) and retention of MAMs (37). To understand the roles of soluble factors in chemotherapy triggered host responses, plasma was collected from GEM treated and control mice. The cytokine/chemokine array data revealed that CCL2 is the most highly expressed chemokine in the plasma of GEM-treated tumor-free mice (Figure 6A). qRT-PCR analysis also showed that CCL2 mRNA expression is increased in the lung tissues of GEM-treated mice (Figure 6B). Previous studies have demonstrated that FXa is able to induce proinflammatory responses in cardiac fibroblasts (38). To assess whether hyper expression of FXa has immune modulatory effect on lung macrophage, we purified lung macrophages from naïve mice and stimulated the cells with recombinant mouse FXa. Cytokine/chemokine array revealed that level of CCL2 is significantly increased in the culture supernatants of FXa-stimulated cells (Figure 6C). Next, we used CCR2 KO mice to determine the importance of CCL2/CCR2 axis in GEM induced pro-metastatic effect. As shown in Figure 6D, deficiency of CCR2 signaling completely abrogated pro-metastatic effect of GEM treatment.

**Figure 6.**
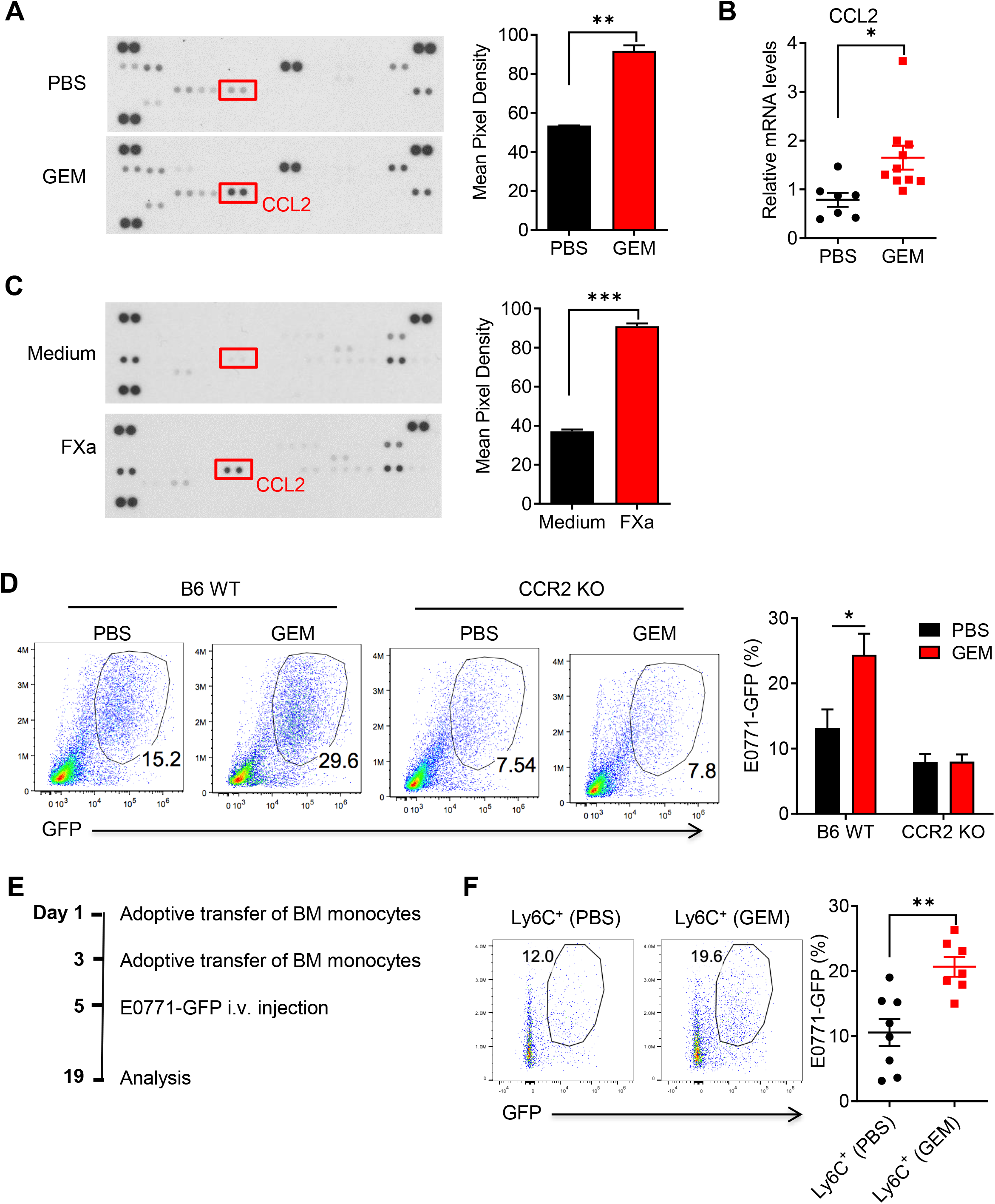
GEM-induced pro-metastatic host response is dependent on CCL2/CCR2 mediated recruitment of monocytes. (**A**) Mouse cytokine/chemokine array of pooled plasma from the mice treated with GEM and PBS. Pixel density of CCL2 was determined using ImageJ. (**B**) qRT-PCR for CCL2 gene in lung tissue from GEM and PBS treated mice (n=7-10). (**C**) Lung macrophages were sorted from naïve mice and stimulated with recombinant mouse FXa (Haematologic Technologies, 1 Unit/ml) for 24 hours. The Mouse cytokine/chemokine profile in the culture supernatant was examined using cytokine/chemokine array. Pixel density of CCL2 was determined using ImageJ. (**D**) B6 WT and CCR2 KO mice were treated with 4 doses of GEM followed by intravenous injection of 4×10^5^ E0771-GFP cells. The percentage of GFP^+^ tumor cells within CD45^-^ cell population were summarized (n=5). (**E**) Experimental scheme for adoptive transfer of BM monocytes (1.5×10^6^/mouse) from PBS- and GEM-treated mice and E0771-GFP (4×10^5^/mouse) injection. (**F**) The lung tumor burden was examined by measuring GFP^+^ tumor cells within lung CD45^-^ cells 14 days later after tumor cell injection (n=7-8). \**p* <0.05, \*\**p* <0.01, \*\*\**p* <0.001.

Functional diversity of monocytes is well recognized because their development can be altered in respond to stimuli (39–42). Our RNAseq data of lung macrophages from GEM vs PBS treated mice also revealed that Proteinase 3 *(Prtn3),* a serine protease enzyme expressed mainly in neutrophils, is highly expressed in lung macrophages from GEM treated mice (Supplemental Fig. 6A). Lung macrophages from GEM-treated mice also demonstrated enrichment of gene signature of neutrophils (Supplemental Figure 6B). We further tested whether lung monocytes from these mice express other neutrophil signature molecules. Indeed, neutrophil elastase *(Elane)* and *S100A8* were increased in lung monocytes from GEM-treated mice (Supplemental Figure 6C). These data suggest that chemotherapy alone induces phenotypic changes of lung monocytes. CCR2 mice are defective in monocyte egress from the BM. To further address the roles of CCR2 and BM monocytes generated via reactive myelopoiesis, BM monocytes were purified from PBS and GEM treated B6 and adoptively transferred into CCR2 KO mice (Figure 6E). Transferred BM monocytes could differentiate into CD11b^+^F4/80^+^ macrophages in lungs (Supplemental Figure 7). We further challenged these mice by intravenous injection of E0771-GFP cells. As shown in Figure 6F, higher tumor burden was observed in the mice transferred with GEM treated BM monocytes compared to the mice transferred with control monocytes. These data suggest that reactive myelopoiesis and CCL2-CCR2 axis play essential roles in GEM induced recruitment of monocytes and macrophages into the lungs, which ultimately contribute into chemotherapy-induced pro-metastatic effect.

### Paclitaxel and Doxorubicin treatment elicits host-responses similar to Gemcitabine

GEM is used in patients with locally advanced or metastatic breast cancer. Having shown that GEM treatment induces pro-metastatic host-responses, we next examined whether similar effects could be triggered by other chemotherapeutic drugs. Paclitaxel (PTX) and Doxorubicin (DOX) are commonly used for the treatment of human breast cancer. Tumor-free naïve mice received a total of three doses of PTX alone or the combination of PTX and DOX and solvent control in two weeks. As shown in Figure 7, PTX and the combination of PTX and DOX treatment induced similar host responses to GEM treatment, including accumulation of lung macrophages (Figure 7A), BM monocytes, LSKs, and MPPs (Figure 7B), and higher expression of FX in lung macrophages (Figure 7C). Importantly, pre-treatment of PTX or the combination of PTX and DOX also induced pro-metastatic effect as revealed by increased lung metastasis in PTX and PTX plus DOX pre-treated mice. Inhibition of FX using FXa inhibitor Rivaroxaban significantly reduced PTX and PTX plus DOX induced pro-metastatic effects (Figure 7D). These findings further support the link between chemotherapy-induced monocyte/macrophage expansion/recruitment, FX expression by these myeloid cells, and FX-induced pro-metastatic effect.

**Figure 7.**
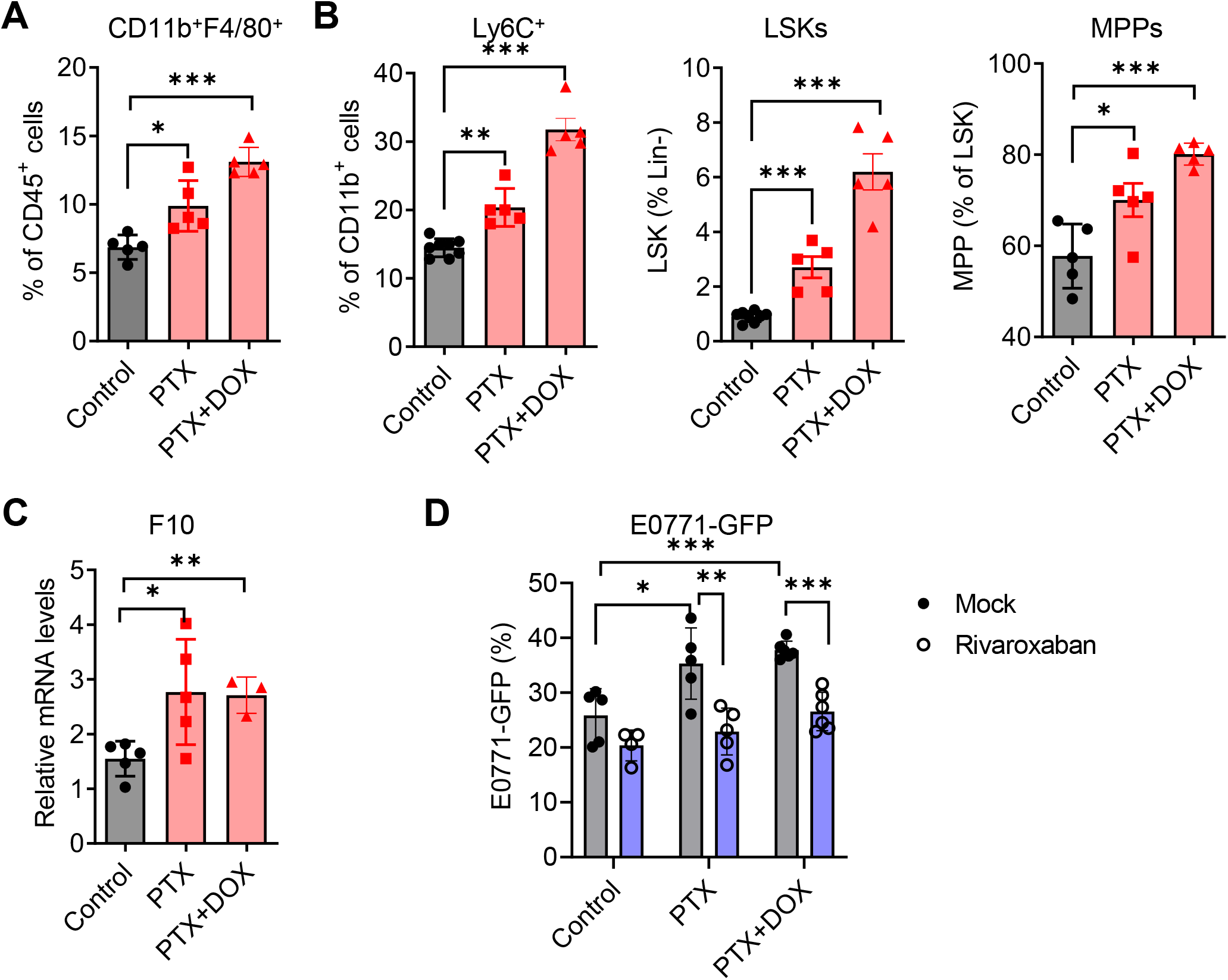
Paclitaxel or combination of Paclitaxel and Doxorubicin treatment elicits host-responses similar to Gemcitabine. B6 WT mice were treated PTX (10 mg/kg, IP), or combination of PTX and DOX (2 mg/kg), or Cremophor EL control 3 times in two weeks. Lung and BM were collected two days later after last treatment. (**A**) Percentage of lung macrophages (CD11b^high^F4/80^+^CD11c^-^) in mice from treated and control mice (n=5). (**B**) Percentages of BM Ly6C^+^ monocytes, LSKs, and MPPs in BM of treated and control mice (n=5-8). (**C**) qRT-PCR for F10 gene in lung macrophages from treated and control mice (n=3-5). (**D**) B6 WT mice were treated with PTX, or combination of PTX and DOX, and Cremophor EL control along with Rivaroxaban or solvent control (mock) for two weeks. E0771-GFP cells (4×10^5^) were intravenously injected into these mice two days later after last treatment. The percentage of GFP^+^ tumor cells in the lung tissues were determined two weeks later (n=5-6). Data are representative of two independent experiments and presented as mean ± SEM. \**p* <0.05, \*\**p* <0.01, \*\*\**p* <0.001.

## Discussion

About 90% of cancer-related deaths are due to metastasis and yet currently there are no broadly effective ways to prevent or cure metastatic cancers (43). Chemotherapy is a mainstay of cancer treatment but in addition to its benefits, chemotherapy can also have the undesired effect of promoting immunosuppression and metastasis (11–13). Previous studies in this area have mainly focused on tumor cell-derived cytokines, chemokines, and exosomes. A recent study showed that chemotherapy elicits pro-metastatic tumor-secreted extracellular vesicles in breast cancer models (14). We also demonstrate that multi-dose chemotherapy enhances the accumulation of immunosuppressive Ly6C^+^ cells in primary tumor microenvironment via the activation of ROS-NF-κB-GM-CSF in tumor cells (18). The current study demonstrates a previously unidentified mechanism by which host responses characterized by accumulation of monocytes/macrophage and up-regulation of macrophage synthesized FX contribute to chemotherapy induced metastasis promotion.

Lung metastasis is the lethal determinant in many cancers. Normal lung tissues constitutively contain abundant myeloid cells which include monocytes, alveolar macrophages, interstitial macrophages (IMs), and neutrophils (4, 7, 44–46). IMs markedly accumulate and differentiate into MAMs in metastatic lung cancer (4). Regarding the origin, lung IMs/MAMs include embryonic tissue-resident macrophages and CCR2-dependent recruited macrophages. Systemic inflammation of primary tumor drives CCR2^+^Ly6C^+^ monocytes migration into lung pre-metastatic niche and differentiation into MAMs (37). We demonstrate that BM derived lung IMs/MAMs are the primary effector cells that mediate pro-metastatic effect of chemotherapy. This conclusion is supported by (1) host-response triggered by chemotherapy promoted monocyte accumulation in the lungs and differentiation into macrophages in the presence or absence of primary tumor, (2) *in vivo* macrophage, neutrophil, and CD4/CD8 T cell depletion studies demonstrated that the pro-metastatic effect of chemotherapy is dependent on macrophages, but not on neutrophils and T cells, (3) the pro-metastatic effect of chemotherapy was abrogated in CCR2 KO mice, and (4) adaptive transfer of BM monocytes into CCR2 KO mice promoted lung metastasis after intravenous injection of tumor cells.

BM suppression is a common side effect of chemotherapy. We observed pro-metastatic of chemotherapy not only at myelosuppression but also at BM recovery phase. Although one time treatment of chemotherapy results in a decrease of MDSC (47, 48), cell depletion induced by cytotoxic drugs might increase myelopoiesis rates to meet the need to replenish depleted reserves (26, 27). This phenomenon is known as reactive myelopoiesis (49). The majority of mobilized HSPCs then differentiate into CD11b^+^Gr-1^+^ cells which limits the efficacy of adoptive T cell therapy (50). In this study, we showed that GEM, PTX, or PTX plus DOX treatment results in myelopoiesis leading to enhanced development of monocytes, but not neutrophils. The difference in myeloid cell development might be due to the different progenitor cells (51, 52). For example, in response to CpG-DNA, monocyte-DC progenitors (MDPs) yielded monocytes and dendritic cells. In contrast, administration of LPS produced neutrophils and monocytes directly from granulocyte-monocyte progenitors (GMP) (52). Further, tumor cells with high metastatic potential enrich MDPs that functionally differentiated into immunosuppressive monocytes to support the metastatic switch (51). Our future study will investigate the effect of chemotherapy treatment on the GMP and MDP. Cytokines/chemokines play important roles in regulating myelopoiesis and HSPC fate decision. In addition to cytokines, ROS also play important roles in regulating HSPCs proliferation and differentiation. In response to inflammation, HSPCs require a metabolic switch from glycolysis to mitochondrial oxidative phosphorylation (OXPHOS) leading an increase of ROS production (53). We showed that GEM treatment increases mtROS production by LSK cells and BM monocytes. This data led us to hypothesize that mtROS might be responsible for GEM induced monocyte development. Indeed, inhibition of mtROS using mitochondria-targeted superoxide scavenger mitoTEMPO abrogates GEM induced hyper differentiation of BM progenitors.

Several molecules, such as VEGFR1 (6), MMP1 (9), TGF-β (54), and IL-35 (55), have been identified to participate in the pro-metastatic effects of lung macrophage. However, current strategies targeting macrophage-associated molecules have shown limited success in clinical settings (3, 56). It’s known that cancer progression results in a higher risk of thromboembolism in cancer patients (57, 58). Recent studies demonstrated that some coagulation factors play important roles in tumor progression and metastasis (57–61). For example, factor XIIIA expressed on lung inflammatory monocytes promoted fibrin cross-linking to create a scaffold for lung squamous cancer cell invasion and metastases (62). FXa treatment promoted tumor and metastasis by inducing endothelial cell activation (59). Additionally, FX expression in myeloid cells within tumor microenvironments resulted in up-regulation of PD-L1 and impaired anti-tumor immunity (63). Some chemotherapy drugs, such as Gemcitabine and Cisplatin, have been associated with an increased risk of thrombotic events (64). Study using factor VIIa inhibitor in combination with GEM showed a non-significant trend toward longer progression-free survival in cancer patients (65). Our study identified FX as one of the highly expressed DEGs in the lung macrophages of tumor-free mice after chemotherapy, suggesting that non-tumor cell-derived factors can induce a hypercoagulable state. Inhibition of FX using oral available FXa inhibitor Rivaroxaban or F10 gene knockdown in macrophages significantly decreased chemotherapy induced pro-metastatic effect. As FX occupies a central position in the coagulation cascade as it can be activated via both the intrinsic and extrinsic pathways (60, 66), our findings suggest that FXa might be a better target for potential combination therapy of chemotherapy and coagulation inhibition.

A limitation of this study is using GEM non-responsible MMTV-PyMT tumor model. Our previous studies have shown that GEM treatment significantly delays the primary tumor progression in E0771 subcutaneous tumor (18). The current study did not observe similar effect in MMTV-PyMT mice. This might be due to the phenotypic difference of two tumor models (22, 67). Non-responsive of GEM on MMTV-PyMT primary tumor progression also could be a net effect of cytotoxic effect of chemotherapy drugs and the pro-tumor effect of reactive myelopoiesis induced systemic immunosuppressive effect. There are two reasons we used MMTV-PyMT model: (1) E0771 is poorly metastatic (68), and (2) It is not comparable to examine the pro-metastatic effect of chemotherapy if the primary tumor sizes are different in chemotherapy-treated and control mice. We understand that use of chemotherapy non-responsive tumor model has limited clinical relevance. But it provides a model to demonstrate pro-metastatic of chemotherapy via the chemotherapy-induced changes in host responses as well as tumor microenvironment.

## Methods

### Mice, tumor models, and chemotherapy in vivo treatment

C57BL/6J mice, CCR2 knockout (KO) mice, and transgenic FVB/MMTV-PyMT mice were purchased from the Jackson Laboratory. Rag2 deficient OVA TCR Tg OT-I mice were purchased from Taconic Biosciences. When maintaining a live colony of FVB/MMTV-PyMT mice, FVB/NJ inbred females were bred with hemizygous males. The pups were genotyped by PCR. Only female transgenic mice were used in studies. All animals were maintained under specific pathogen-free conditions and handled in accordance with the protocols approved by the Institutional Animal Care and Use Committee of the University of Louisville (IACUC 21873). The murine mammary cancer cell lines E0771 (from CH3 BioSystems) and E0771-GFP cells (Yan Lab) were confirmed pathogen free by VRL Diagnostics without further authentication. Cells were cultured in the completed DMEM medium containing 10% FBS and used at < 16 passages. To establish subcutaneous (s.c.) tumor, 1×10^6^ E0771 tumor cells were suspended in PBS and inoculated s.c. in the flank of female B6 mice. Tumor volumes were calculated using the formula V=(width*width*length)/2. For GEM *in vivo* treatment, 9-10 weeks old transgenic female MMTV-PyMT, E0771 tumor-bearing mice (tumor size between 6-8 mm), or naïve (tumor-free) mice were treated with GEM (60 mg/kg body weight, Accord Healthcare Inc.) or PBS by intraperitoneal (IP) injection 4 times in two weeks. For PTX and DOX *in vivo* treatment, tumor-free mice were treated with PTX (10 mg/kg, IP, Sigma, Cat. No. T7402), DOX (2 mg/kg, IV, TOCRIS, Cat. No. 2252), or Cremophor EL control 3 times in two weeks. The mice were sacrificed 2 days later after the last treatment to analyze cellularity within BM and lungs. For experimental metastasis model, E0771-GFP cells (4×10^5^) were intravenously injected into B6 WT, Ly6G or CD4/CD8 depletion antibody treated, or CCR2 KO mice pre-treated with chemotherapy. The mice were euthanized two weeks later after tumor cell injection. To quantify metastatic tumor burden, GFP^+^ tumor cells were quantified by Flow cytometry. Lung tissue sections were stained with H&E and scanned with 3DHISTECH’s CaseViewer.

### Preparation of lung single cell suspension, flow cytometry, and cell sorting

Lungs were washed extensively in PBS, and then minced before digestion with collagenase IV (0.2 mg/ml), dispase (2 mg/ml) and DNase I (0.002 mg/ml) in complete RPMI1640 medium for 30 min at 37°C. After incubation, the digestion was immediately stopped by addition of 5 ml cold medium. The suspension was then filtered through a 40 μm cell strainer into petri dishes and extra tissue chunks were further mashed with syringe columns. The red blood cells were lysed by adding ACK lysis buffer for about 1 min and washed twice with complete medium. For flow cytometry analysis, the cells were blocked in the presence of anti-CD16/CD32 at 4°C for 10 min and stained on ice with the appropriate antibodies and isotype controls in PBS containing 1% FBS. The fluorochrome-labeled antibodies against mouse CD45, CD11b, Ly6G, Ly6C, F4/80, CD4, CD8, IFN-γ, granzyme B, Foxp3 and their corresponding isotype controls were purchased from BioLegend. APC-conjugated anti-mouse CCR2 Ab was from R&D Systems. Fixable Viability Dye eFluor™ 780 was from Thermo Fisher Scientific. For intracellular staining, the cells were fixed and permeabilized following surface staining. The samples were acquired using FACSCanto cytometer (BD Bioscience) or Cytek Aurora cytometry and analyzed using FlowJo software. The lung macrophage population (CD45^+^CD11b^high^F4/80^+^CD11c^-^) was sorted by using BD FACS Aria III. Fixable viability dye was used to exclude dead cells. A post sort analysis was performed to determine the purity of the macrophages with approximately 90% purity. Antibodies used in flow cytometry were listed in Supplemental Table 1.

### Reactive myelopoiesis analysis

BM suspensions were prepared by harvesting mouse femur and tibia bones and flushing them with complete medium. The cells were harvested, and the red blood cells were lysed by using ACK lysis buffer. The cells were washed twice and suspended in complete medium. For BM stem and progenitor analysis, BM cells were stained with Abs for lineage markers (CD19, Ter119, CD11b, Ly6G/C, CD3, NK1.1) along with anti-Ly6A/E (Sca-1), anti-CD117 (c-kit), anti-CD48 and anti-CD150 (BioLegend) for Lin^-^Sca-1^+^c-kit^+^ (LSK) cell population and multipotent progenitors (MPPs). For BM cell *in vitro* differentiation assay, BM cells were cultured in the presence of recombinant mouse GM-CSF (20 ng/ml) (35) or 20% E0771 conditioned medium (CM) for 2 days. The culture medium including non-adherent cells was entirely discarded at day 3 and replaced by medium containing GM-CSF or E0771 CM for additional four days. The phenotype of *in vitro* cultured cells was analyzed by flow cytometry. Immunosuppression assay of BM Ly6C^+^ cells was performed by co-culturing with CFSE labeled OT-I splenocytes in the presence of OVA (18). For assessment of granulocyte/monocyte colony forming unit (CFU-GM) colony, BM cells (5×10^3^) were seeded in MethoCult™ GF M3534 (STEM Technologies, Cat. No. 03534) methylcellulose-based medium containing recombinant cytokines for mouse myeloid progenitor cells. The colonies of CFU-M and CFU-GM with more than 50 cells were determined after 7 days of culture.

### CyTOF mass cytometry data acquisition and analysis

Lung cell staining was performed according to the protocol of Maxpar Cell Surface Staining with Fresh Fix (Fluidigm). Prior to acquisition, cells were suspended in a 1:9 solution of cell acquisition solution: EQ 4 element calibration beads (Fluidigm) at an appropriate concentration at no more than 600 events per second. Data acquisition was performed on the CyTOF Helios system (Fluidigm). FCS files were normalized with Helios instrument work platform (FCS Processing) based on the calibration bead signal used to correct any variation in detector sensitivity. CyTOF data analysis was performed with FlowJo software. Total events were gated after removing beads, doublets, and dead cells (uptake of cisplatin). tSNE and FlowSOM clustering analysis for CyTOF data were performed using FlowJo Plugins platform. tSNE analysis was performed on all samples combined. Different immune populations were defined by the expression of specific surface markers. Antibodies used in CyTOF were listed in Supplemental Table 2.

### Cytokine array and ELISA

The cytokine/chemokine profile of mice plasma or culture supernatants was determined by using proteome profiler mouse cytokine array kit (R&D Systems, CAT. No. ARY006). The expression levels of cytokine/chemokine were measured by pixel density of each dot using the ImageJ software. Mouse Factor X total antigen was measured using kit from Molecular Innovations (Cat. No. MFXKT-TOT).

### Quantitative real-time PCR

Small pieces of lung tissue or sorted lung macrophages were frozen in TRIzol (Invitrogen) at −80°C. RNA was extracted and transcribed to cDNA with a reverse transcription kit (Bio-Rad). Quantitative real-time PCR reactions were performed using SYBR Green Supermix (Bio-Rad). We normalized gene expression level to ribosomal protein L13a (RPL13A) housekeeping gene and represented data as fold differences by the 2^-ΔΔCt^ method. The primer sequences of real-time PCR were as follows: F10: F: 5’-GACAATGAAGGGTTCTGTGG-3’ and R: 5’-CTGTGTTCCGATCACCTACC-3’; CCL2: F: 5’-GGTCCCTGTCATGCTTCTGG-3’ and R:5’-GCTGCTGGTGATCCTCTTGT-3’; S100A8: F: 5’-GGAGTTCCTTGCGATGGTGAT-3’ and R: 5’-GTAGACATATCCAGGGACCCAGC-3’; S100A9: F: 5’-AGCATAACCACCATCATCGACAC-3’ and R: 5’-TGTGCTTCCACCATTTGTCTGA-3’; TGF-β: F:5’-TGCTAATGGTGGACCGCAA-3’ and R: 5’-CACTGCTTCCCGAATGTCTGA-3’. Each sample was run in duplicate.

### Mitochondrial reactive oxygen species (mtROS), mitochondrial potential, and dysfunctional mitochondria measurement

For quantification of mitochondrial ROS levels, BM cells were incubated with MitoSOX Red (5 μM, Thermo Fisher Scientific, Cat. No. M36008) at 37 °C in PBS and then with appropriate BM progenitor or monocyte makers plus viability dye for 20 min at 4 °C. The levels of mtROS were determined by Flow cytometry. The mitochondrial membrane potential and dysfunctional mitochondria were measured by staining BM cells with Tetramethylrhodamine Methyl Ester (TMRM 500 nM, Thermo Fisher Scientific, Cat. No. T668), MitoTracker Green (100 nM, Thermo Fisher Scientific, Cat. No. M7514), and MitoTracker Red (50 nM, Thermo Fisher Scientific, Cat. No. M7512) for 30 min at 37 °C in PBS and then with appropriate BM progenitor makers plus viability dye for 20 min at 4 °C.

### RNA sequencing and analysis

Lung macrophages from PBS and GEM-treated tumor-free mice were sorted, and RNA was extracted using a QIAGEN RNAeasy Kit (QIAGEN). The quantity of the purified RNA samples was measured by the RNA High Sensitivity kit in the Qubit Fluorometric Quantification system (Thermo Fisher Scientific). Libraries were prepared using the Universal Plus mRNA-Seq with NuQuant (NuGEN). Pooled library was run on MiSeq to test quantity and quality, using the MiSeq Reagent Nano Kit V2 300 cycles (Illumina). Sequencing was performed on the Illumina NextSeq 500 using the NextSeq 500/550 75 cycle High Output Kit v2.5. Differential expression was performed using two different tools, DESeq2 and Cuffdiff2. Differentially expressed genes (DEGs) at p-value cutoff 0.01, q-value cutoff of 0.01 with log2FC of 0 for the pairwise comparisons were used for further analysis of enriched Gene Ontology Biological Processes (GO:BP). Volcano plot was created to examine the distribution of log2 fold change at different significance levels. RNA-seq data were deposited with GEO accession (GSE217105).

### Rivaroxaban in vivo treatment

Rivaroxaban (Sigma-Aldrich, Cat. No. SML2844) is an orally active, direct inhibitor of Factor Xa. A stock solution was made by dissolving the Rivaroxaban in DMSO. After mixing with 0.5% methylcellulose + 0.2% Tween 80, Rivaroxaban (20 mg/kg) or solvent control was orally administered to mice daily via oral gavage for 2 weeks.

### Neutrophil, T cell, and macrophage in vivo depletion

Depletion of neutrophils, CD4 and CD8 T cells was performed by IP injection of anti-Ly6G mAb (300 μg, twice a week, Bio X Cell, Cat. No. BE0075-1), anti-CD4 (clone GK1.5, Yan Lab) and anti-CD8 mAb (clone 2.43, Yan Lab, weekly, 250 μg). Macrophage depletion was performed by IV injection of Clodrosome (200 μl/mouse, 3 times per week, Encapsula NanoSciences, Cat. No. CLD-8909). The depletion efficiency was examined by Flow cytometry.

### Adoptive transfer of BM monocytes and BM-derived macrophages

BM cells were harvested from PBS and GEM-treated tumor-free mice. BM monocytes were purified using mouse BM monocyte isolation kit (Miltenyi Biotec, Cat. No. 130-100-629) and AutoMACS Pro Separator. The purified monocytes (1.5×10^6^) were adoptively transferred into CCR2 KO mice twice with 48 hours apart. For macrophage F10 gene knockdown, BM-derived macrophages were transfected with F10 siRNA (Thermo Fisher Scientific, Cat. No. AM16706) and control siRNA (Thermo Fisher Scientific, Cat. No. AM4635) and Lipofectamine™ RNAiMAX transfection reagent (Thermo Fisher Scientific, Cat. No. 13778030) followed by stimulation with 20% E0771 conditioned medium (CM) for 24 hours. The control and F10 gene knockdown macrophages (2×10^6^) were adoptively transferred into CCR2 KO mice twice with 48 hours apart. E0771-GFP cells were IV injected into CCR2 KO recipient mice 48 hours later and lung metastasis was determined 14-17 days later after tumor cell injection.

### Statistics

Data were analyzed using GraphPad Prism software (GraphPad Software). An unpaired Student t test and one-way or two-way ANOVA were used to calculate significance. All graph bars are expressed as mean ± SEM. Significance was assumed to be reached at p < 0.05. The p values were presented as follows: *p<0.05, **p<0.01, and ***p<0.001.

## Author contributions

CW, QZ, JY, and CD conceptualized and designed the study. CW, QZ, RS, JW, XH, and HL developed the methodology and acquired data. ECR analyzed RNAseq data. CW, QZ, JY, and CD analyzed and interpreted data. JY and CD wrote manuscript and supervised study.

## Acknowledgments

The authors wish to thank Sabine Waigel from the Brown Cancer Center Genomics Facility. This work was supported by the NIH P20GM135004. Part of this work was performed with assistance of the UofL Genomics Facility, which is supported by NIH P20GM103436 (KY IDeA Networks of Biomedical Research Excellence)

