## Supplemental data for "Reactive myelopoiesis and FX-expressing monocyte-derived macrophages triggered by chemotherapy promote cancer lung metastasis"

**Supplemental Table 1. Antibodies used in Flow cytometry and Western blotting**

| Antibodies | Company and catalog number |
|---|---|
| PerCP/Cyanine5.5 anti-mouse CD45 | BioLegend No. 157208 |
| PE/Cyanine7 anti-mouse CD45 | BioLegend No. 103114 |
| APC anti-mouse/human CD11b | BioLegend No. 101212 |
| PE anti-mouse Ly6G | BioLegend No. 127608 |
| PerCP anti-mouse Ly6C | BioLegend No. 128028 |
| PE anti-mouse F4/80 | BioLegend No. 123110 |
| FITC anti-mouse CD11c | BioLegend No. 117306 |
| APC anti-mouse CD3 | BioLegend No. 100236 |
| APC anti-mouse CD4 | BioLegend No. 100412 |
| FITC anti-mouse CD8a | BioLegend No. 100706 |
| APC anti-mouse NK-1.1 | BioLegend No. 108710 |
| APC anti-mouse CD19 | BioLegend No. 115512 |
| APC anti-mouse Ly-6G/Ly-6C | BioLegend No. 108412 |
| APC anti-mouse TER-119 | BioLegend No. 116212 |
| Anti-mouse Ly6A/E (Sca-1) | BD Bioscience No. 553108 |
| PE/Cyanine7 anti-mouse CD117 (c-kit) | BioLegend No. 135112 |
| FITC anti-mouse CD48 | BioLegend No. 103404 |
| PE/Cyanine5 anti-mouse CD150 (SLAM) | BioLegend No. 115912 |
| PE anti-mouse IFN-γ | BioLegend No. 505808 |
| PE anti-human/mouse Granzyme B | BioLegend No. 372208 |
| PE anti-mouse FoxP3 | BioLegend No. 126404 |
| APC-conjugated anti-mouse CCR2 | R&D Systems No. FAB5538A-100 |
| Fixable Viability Dye eFluor™ 780 | Thermo Fisher Scientific No. 65-0865-14 |
| Polyclonal Rabbit anti-Human F10 / Factor X Antibody | LifeSpan BioSciences No. LS-C331476 |

**Supplemental Table 2. Antibodies used in CyTOF**

| | **Antibodies** | **Company and catalog number** |
|---|---|---|
| 1 | Anti-Mouse CD45 (30-F11)-89Y | Fluidigm, No. 3089005B |
| 2 | Anti-Mouse Ly-6G (1A8)-141Pr | Fluidigm, No. 3141008B |
| 3 | Anti-Mouse CD11c (N418)-142Nd | Fluidigm, No. 3142003B |
| 4 | Anti-Mouse CD69 (H1.2F3)-143Nd | Fluidigm, No. 3143004B |
| 5 | Anti-Mouse CD4 (RM4-5)-145Nd | Fluidigm, No. 3145002B |
| 6 | Anti-Mouse F4/80 (BM8)-146Nd | Fluidigm, No. 3146008B |
| 7 | Anti-Mouse CD103 (2E7)/148Nd | Biolegend, No. 121402 |
| 8 | Anti-Mouse CD19 (6D5)-149Sm | Fluidigm, No. 3149002B |
| 9 | Anti-Mouse Ly-6C (HK1.4)-150Nd | Fluidigm, No. 3150010B |
| 10 | Anti-Mouse CD25 (3C7)-151Eu | Fluidigm, No. 3151007B |
| 11 | Anti-Mouse CD3e (145-2C11)-152Sm | Fluidigm, No. 3152004B |
| 12 | Anti-Mouse CD274/PD-L1-153Eu (10F.9G2) | Fluidigm, No. 3153016B |
| 13 | Anti-Mouse PD-1 (29F.1A12)-159Tb | Fluidigm, No. 3159024B |
| 14 | Anti-Mouse CD62L (MEL-14)-160Gd | Fluidigm, No. 3160008B |
| 15 | Anti-Human/Mouse CD44 (IM7)-162Dy | Fluidigm, No. 3162030B |
| 16 | Anti-Mouse CX3CR1 (SA011F11)-164Dy | Fluidigm, No. 3164023B |
| 17 | Anti-Mouse CD8a (53-6.7)-168Er | Fluidigm, No. 3168003B |
| 18 | Anti-Mouse CD206/MMR (C068C2)-169Tm | Fluidigm, No. 3169021B |
| 19 | Anti-Mouse NK1.1 (PK136)-170Er | Fluidigm, No. 3170002B |
| 20 | Anti-Mouse CD11b (M1/70 )-172Yb | Fluidigm, No. 3172012B |
| 21 | Anti-Mouse CD223/LAG3 (C9B7W)-174Yb | Fluidigm, No. 3174019B |
| 22 | Anti-Human/Mouse CD45R/B220 (RA3-6B2)-176Yb | Fluidigm, No. 3176002B |
| 23 | Anti-Mouse I-A/I-E (M5/114.15.2)-209Bi | Fluidigm, No. 3209006B |
| 24 | Anti-Mouse CD127/IL7Ra (A7R34)-175Lu | Fluidigm, No. 3175006B |

**Supplemental Figure 1**

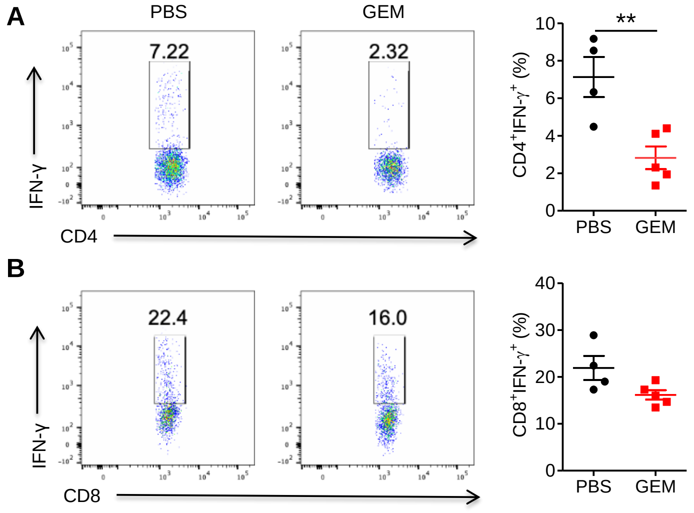


**Figure S1.** Mice bearing subcutaneous E0771 tumor cells were treated with 4 times of GEM in two weeks. Lung tissues were collected 2 days later after last GEM treatment. Cells were stimulated with PMA/Ionomycin in the presence of protein transport inhibitor brefeldin A for 4 hours. IFN-γ producing CD4 T cells (**A**) and CD8 T cells (**B**) were determined by intracellular cytokine staining and Flow cytometry. Each dot represents one mouse (n=4-5). ***p* <0.01.

**Supplemental Figure 2**

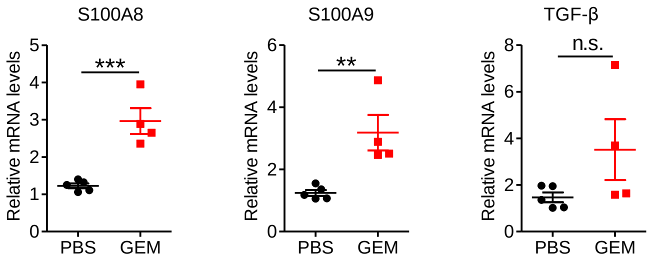


**Figure S2.** Naïve mice were treated with 4 times of GEM in two weeks. Lung tissues were collected 48 hours later after last treatment. Gene expression of S100A8, S100A9, and TGF-β in lung tissues from GEM and PBS treated mice was determined with qRT-PCR. Each dot represents one mouse (n=4-5). ***p* <0.01, ****p* <0.001.

### Supplemental Figure 3

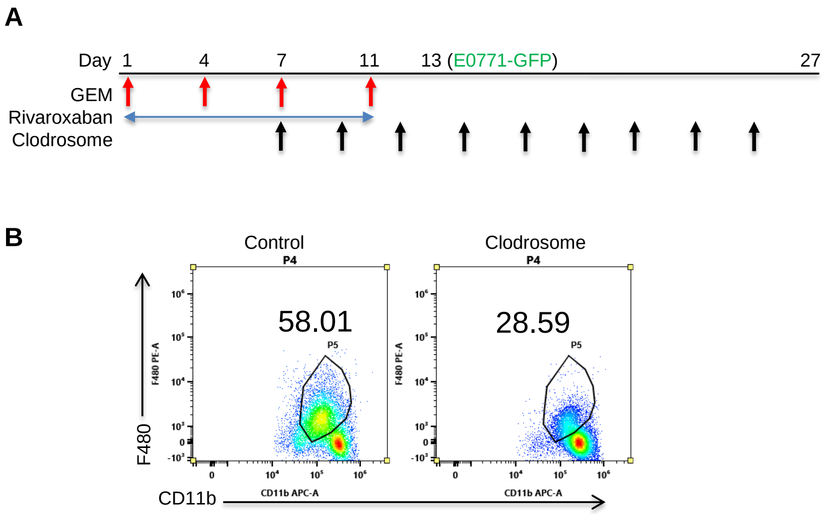


**Figure S3.** (**A**) Schema for *in vivo* GEM and Rivaroxaban treatment, macrophage depletion, and E0771-GFP tumor cell intravenous injection. (**B**) Lung macrophage depletion efficacy after intravenous injection of 3 times of Clodrosome was determined by staining cells with CD45, CD11b, F4/80, and viability dye. Cells were gated on $CD45^{+}CD11b^{+}$ population.

### Supplemental Figure 4

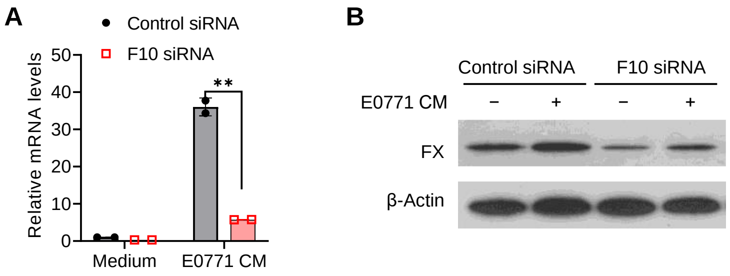


**Figure S4.** BM-derived macrophages were transfected with control siRNA and F10 siRNA. 24 hours later cells were treated with 20% E0771 conditioned medium (CM) for 24 hours. F10 gene knock down efficiency was determined by using qRT-PCR (**A**) and Western blotting (**B**).

### Supplemental Figure 5

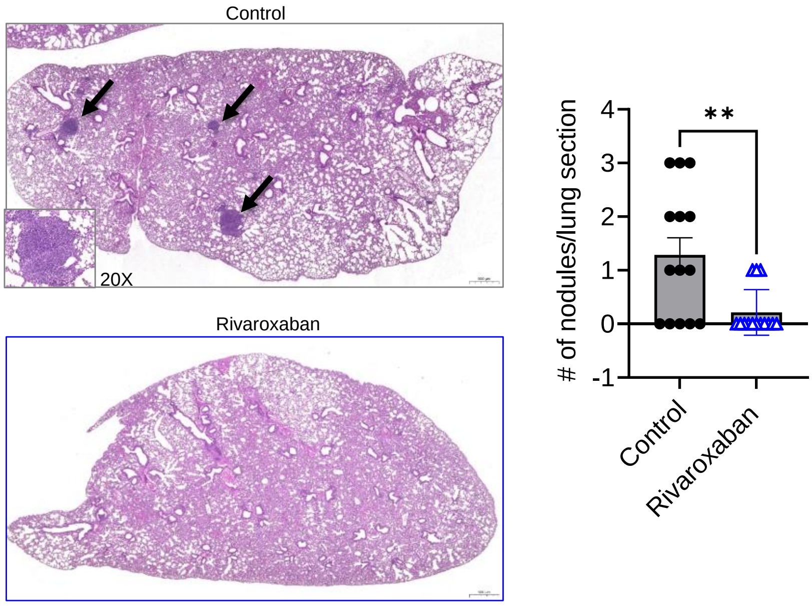


**Figure S5.** E0771-GFP subcutaneous tumor-bearing mice (primary tumor size between 6-8 mm in diameter) were treated with Rivaroxaban (oral, 20 mg/kg, daily) or solvent control for two weeks. Lung tissues were collected at day 31 after tumor cell injection. The metastasis in lungs was determined by evaluation of tumor nodules using H&E staining. One section represents one mouse (n=14). Numbers of nodule per lung section were counted and summarized. ***p* <0.01.

**Supplemental Figure. 6**

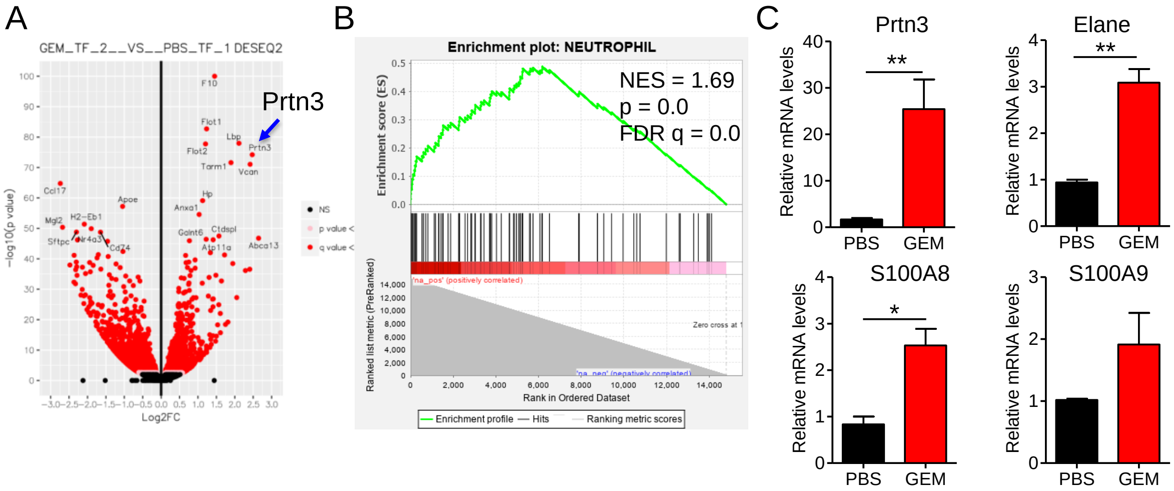


**Figure S6.** (**A**) Differential gene expression in sorted macrophages from GEM-treated tumor-free mice as compared to macrophages from PBS-treated mice. Gene *Prtn3* is one of DEGs in lung macrophages. (**B**) Analysis of lung macrophage RNA-seq data revealed the enrichment plot with upregulation of neutrophil signature genes in lung macrophages from GEM-treated versus PBS-treated tumor-free mice. (**C**) Lung monocytes ($CD45^{+}CD11b^{+}Ly6C^{high}$) were sorted for gene expression analysis using qRT-PCR. **p* <0.05, ***p* <0.01.

**Supplemental Figure. 7**

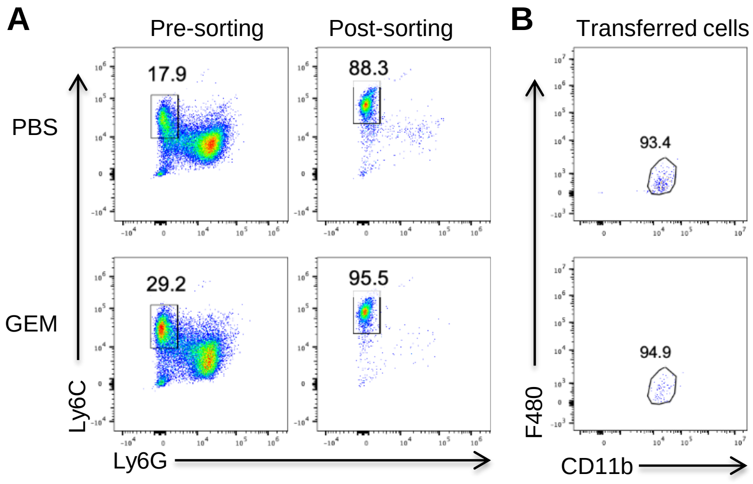


**Figure S7.** (**A**) Percentages of BM monocytes from PBS and GEM treated mice before and after purification. (**B**) CFSE labeled purified monocytes (1.5x$10^6$) were intravenously injected into CCR2 KO mice. Expression of CD11b and F480 in transferred cells (CFSE$^+$) in the lungs analyzed 48 hours later.
